# SPAC: a scalable, integrated enterprise platform for end-to-end single cell spatial analysis of multiplexed tissue imaging

**DOI:** 10.1101/2025.04.02.646782

**Authors:** Fang Liu, Rui He, Thomas Sheeley, David Scheiblin, Stephen J Lockett, Lisa A Ridnour, David A Wink, Mark Jensen, Janelle Cortner, George Zaki

## Abstract

**Background:** Multiplexed tissue imaging enables the simultaneous detection of dozens of proteins at single-cell resolution, providing unprecedented insights into tissue organization and disease microenvironments. However, the resulting high-dimensional, gigabyte-scale datasets pose significant computational and methodological challenges. Existing analytical workflows, often fragmented between bespoke scripts and static visualizations, lack the scalability and user-friendly interfaces required for efficient, reproducible analysis. To overcome these limitations, we developed SPAC (analysis of SPAtial single-Cell datasets), a scalable, web-based ecosystem that integrates modular pipelines, high-performance computing (HPC) connectivity, and interactive visualization to democratize end-to-end single-cell spatial analysis applied to cellular positional data and protein expression levels.

**Results:** SPAC is built on a modular, layered architecture that leverages community-based and newly developed tools for single-cell and spatial proteomics analysis. A specialized Python package extends these functionalities with custom analysis routines and established software engineering practices. An Interactive Analysis Layer provides web-hosted pipelines for configuring and executing complex workflows, and scalability enhancements that support distributed or parallel execution on GPU-enabled clusters. A Real-Time Visualization Layer delivers dynamic dashboards for immediate data exploration and sharing. As a showcase of its capabilities, SPAC was applied to a 4T1 breast cancer model, analyzing a multiplex imaging dataset comprising 2.6 million cells. GPU acceleration reduced unsupervised clustering runtimes from several hours to under ten minutes, and real-time visualization enabled detailed spatial characterization of tumor subregions.

**Conclusions:** SPAC effectively overcomes key challenges in spatial single-cell analysis by streamlining high-throughput data processing and spatial profiling within an accessible and scalable framework. Its robust architecture, interactive interface and ease of access have the potential to accelerate biomedical research and clinical applications by converting complex imaging data into actionable biological and clinical insights.

## 1. Background

Multiplexed tissue imaging technologies now enable the simultaneous detection of dozens of proteins at single-cell resolution within a single tissue section. Techniques such as imaging mass cytometry (IMC), multiplexed ion beam imaging (MIBI), multiplexed immunofluorescence (MxIF), cyclic immunofluorescence (CyCIF), co-detection by indexing (CODEX), and others (e.g., 4i, mIHC, IBEX, COMET, InSituPlex, and Vectra) allow whole-slide staining and capture spatially resolved protein expression patterns [1–11]. Computational pipelines including HALO [12] and MCMICRO [13] leverage segmentation and pixel-level deep learning to delineate cell boundaries and extract per-cell features (e.g., staining intensities, morphological measurements, and spatial coordinates). This wealth of spatial information opens new avenues for understanding tissue organization and disease microenvironments but also introduces significant methodological and computational challenges.

A central challenge lies in translating high-dimensional, spatially resolved protein data into biologically meaningful insights. Critical tasks include identifying distinct cell subsets, characterizing cell-cell interactions, and detecting spatial patterns that correlate with clinical or experimental conditions. Although tools such as Seurat [14], GraphST [15], and Bento [16] have advanced single cell spatial transcriptomics analysis, their workflows and data structures are not tailored to high-plex image protein data. Similarly, while SPIAT [17], Giotto [18], Squidpy [19] and SCIMAP [20] can be applied to both spatial transcriptomic and spatial proteomic dataset, they typically lack a user-friendly graphical interface capable of handling the large-scale, image-derived single-cell data generated by these technologies.

Moreover, the traditional division of labor in research teams further hampers efficient data analysis. Bench scientists are often responsible for hypothesis generation, experimental design, and data production, while data scientists (e.g., bioinformaticians and image analysts) develop and maintain custom scripts or Jupyter notebooks to analyze these complex datasets. This separation frequently results in slow, fragmented workflows, where data are exchanged across disparate environments, analyses are performed on specialized computational setups, and results are delivered as static visual outputs (e.g., PowerPoint slides). Such compartmentalization hinders iterative exploration, slows hypothesis testing, and can compromise reproducibility.

High-resolution whole-slide imaging (WSI) compounds these challenges by producing datasets that may reach hundreds of gigabytes in size, encompassing millions of cells over extensive tissue areas (e.g., a 50-plex, 4 cm² slide at 0.3 μm resolution). Under these conditions, computation-intensive tasks become bottlenecks on standard workstations. For instance, graph-based clustering methods such as PhenoGraph [21] construct k-nearest neighbor graphs based on Euclidean distances using normalized cellular features, refine these graphs by adjusting edge weights according to the number of shared neighbors between cells, and then apply community detection algorithms (e.g., Leiden or Louvain) to delineate clusters. When combined with iterative clustering across multiple parameters, and dimensionality reduction techniques such as UMAP [22], these methods can require tens of gigabytes of memory for datasets containing over 10 million cells and tens of markers. Consequently, there is a pressing need for platforms that combine user-friendly interfaces, scalable pipelines, and direct integration with HPC or cloud-based resources.

SPAC (analysis of SPAtial single-Cell datasets) overcomes these limitations by leveraging a scalable enterprise platform that integrates open-source Python packages with an interactive analysis layer and a real-time visualization module (Fig. 1). Its overarching goal is to democratize spatial single-cell workflows through several dimensions:

- **Browser-Based Access:** Researchers can perform state-of-the-art analyses directly within a web browser, eliminating the need for local installations. Bench scientists can easily launch workflows, adjust parameters, and visualize results interactively—without relying on command-line expertise.
- **Integrated Pipeline:** SPAC offers a modular yet unified environment that encompasses exploratory data analysis (EDA), quality control, clustering, phenotyping, and spatial analysis. This cohesive pipeline maintains a consistent data lineage, minimizes burdens associated with data format conversion, and ensures reproducibility by preserving a detailed, shareable record of all computational steps and parameters—even as personnel change.
- **Scalability and HPC Integration:** SPAC seamlessly connects to on-premise or cloud-based HPC resources, providing compute nodes over 150 GB of memory and specialized accelerators (e.g.,GPUs). Automated load balancing, parallel computing, and containerized execution empower users of all levels to process large-scale datasets containing millions of cells rapidly and efficiently.
- **Collaboration and Reproducibility:** SPAC’s architecture fosters a collaborative research environment where biologists can pose hypotheses and interact with dynamic visualizations (such as spatial maps, heatmaps, boxplots), while data scientists refine and configure robust analytical pipelines. Integrated features like version control, parameter tracking, and shared project workspaces ensure that analyses remain reproducible and transparent.

**Fig. 1.**
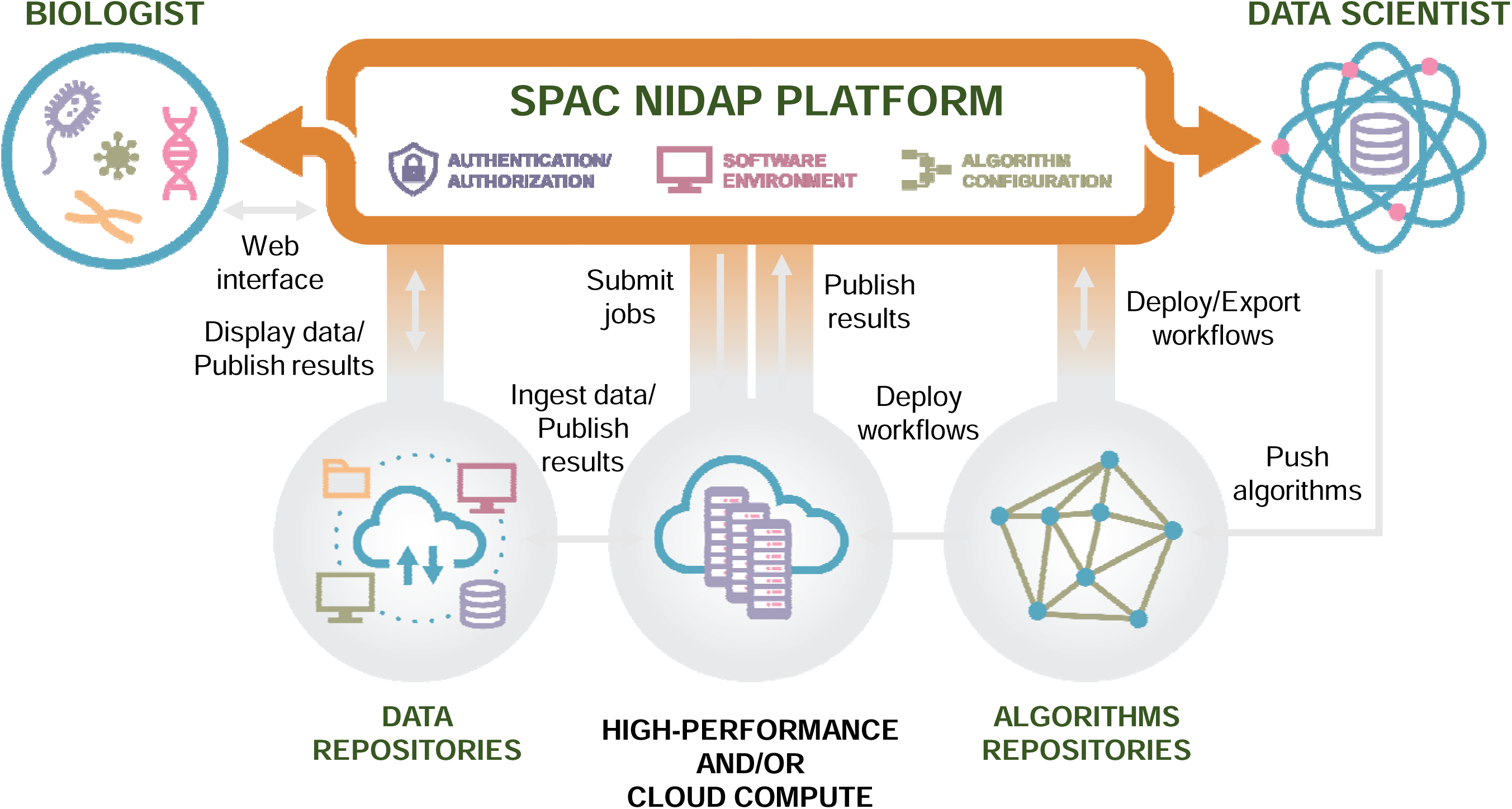
Overview of the SPAC Ecosystem. SPAC integrates with the NIDAP platform to support both bench biologists and data scientists. Biologists upload imaging data (e.g., CSV outputs from HALO, MCMICRO, QuPath, Visiopharm) into NIDAP, where SPAC converts per-cell tables into AnnData format. Compute-intensive tasks, such as clustering, dimensionality reduction, and spatial statistics, are offloaded to HPC resources via the NIDAP HPC Connector, eliminating the need for direct HPC logins. Data scientists deploy new algorithms and workflows from SPAC’s repositories. SPAC’s interactive pipelines streamline analysis from data sampling and preprocessing to clustering and spatial analysis and automatically return results to NIDAP, enabling web-based exploration of large-scale, HPC-powered analyses with transparent monitoring and shareable reports.

In summary, SPAC unites the strengths of existing spatial omics tools with the convenience and scalability demanded by advanced studies of complex tissues and disease microenvironments. By bridging the gap between biologists and computational experts, SPAC accelerates the transformation of high-dimensional, cell-segmented multiplexed imaging data into actionable biological insights.

## 2. Implementation

### 2.1 SPAC Ecosystem

The SPAC ecosystem is designed with a modular, layered architecture that leverages community-standard tools while providing enterprise-level scalability, interactive pipelines, and real-time visualization (Fig. 2). Each layer in the stack is tailored to accommodate different user groups from software engineers and data scientists developing new workflows to bench scientists and principal investigators who primarily review and interpret results without installing or maintaining complex software.

**Fig. 2.**
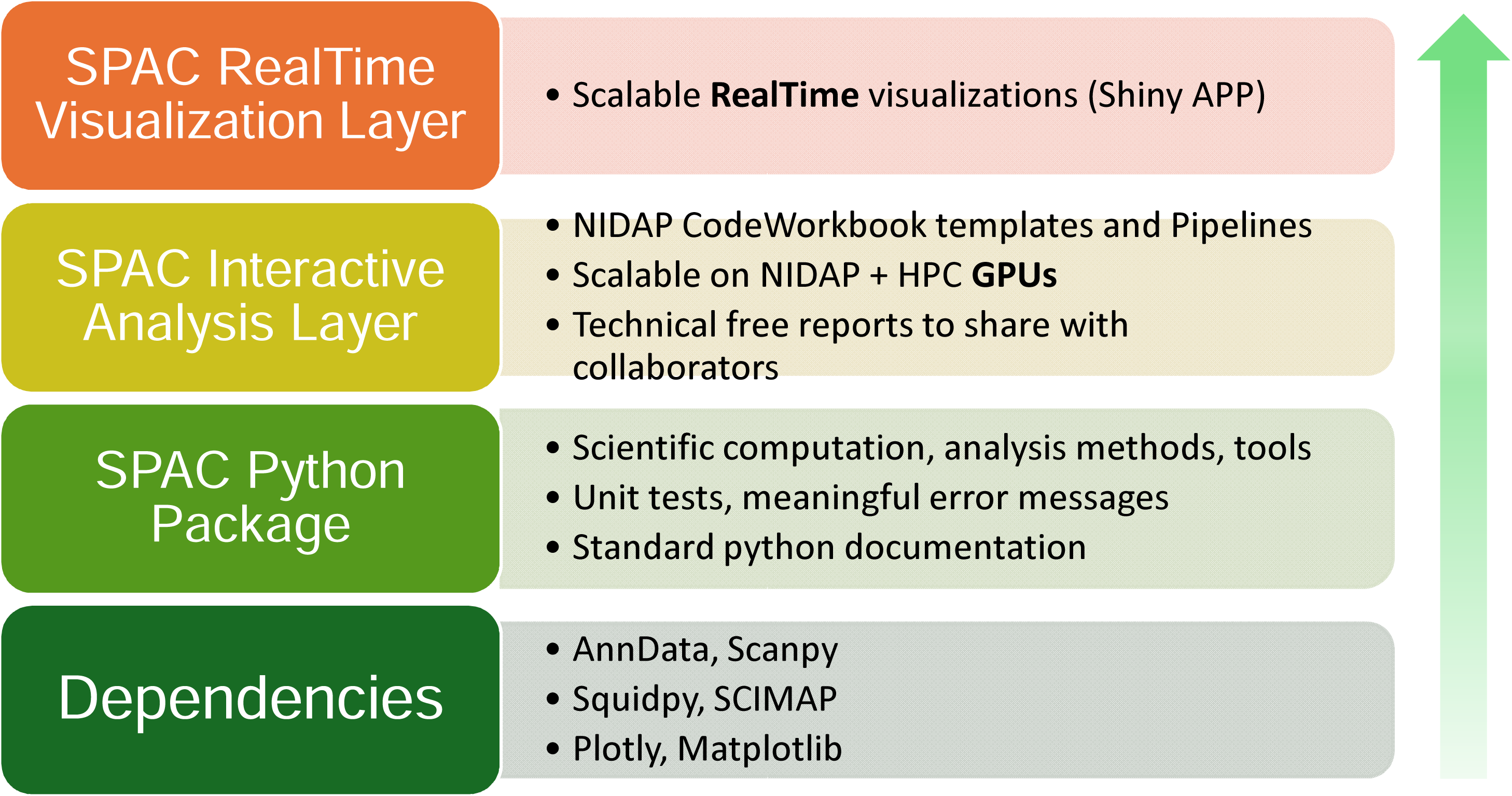
The SPAC Ecosystem: A Modular Framework for Scalable Imaging Analysis. SPAC has a four-layer architecture. The Foundational Dependencies Layer integrates open-source tools for core single-cell and spatial proteomics methods. The SPAC Python Package Layer adds custom analysis routines, rigorous software practices, and GPU-enabled HPC support. The Interactive Analysis Layer delivers web-based pipelines via NIDAP Code Workbook templates, scaling automatically to HPC resources and generating shareable reports. The Real-Time Visualization Layer decouples heavy computations from the front end, enabling dynamic exploration of large spatial datasets through dashboards and live annotations.

#### 2.1.1 Foundational Dependencies

At the base of the SPAC stack lie established open-source packages for single-cell and spatial proteomics analysis, including AnnData [23], SCANPY [24], Squidpy, and SCIMAP. These tools, integral to the “scverse” [25] ecosystem, provide data structures and methods that have become de facto standards in Python-based single-cell analytics. The scverse ecosystem is a collaborative community-driven initiative that unifies diverse single-cell analysis tools into a coherent framework, promoting interoperability, reproducibility, and standardized workflows. By building on these foundational libraries, SPAC aligns with community best practices, ensures ongoing compatibility with widely adopted formats, and inherits robust core functionalities such as dimensionality reduction, clustering, spatial graph construction.

#### 2.1.2 SPAC Python Package

Directly built on top of these community tools, the SPAC Python Package extends and augments existing functionalities to meet the specific needs of multiplexed tissue imaging workflows. It implements specialized analysis routines that customizes preprocessing input dataset (e.g., batch correction, normalization and data transformation), sample stratification, phenotyping, HPC-enabled clustering schemes, and spatial statistics that are not available in standard packages. SPAC also supports exporting processed data, enabling researchers to perform independent downstream analyses using external tools. In parallel, the package adheres to rigorous research software engineering principles by incorporating comprehensive unit tests, robust handling of edge cases, informative error messages, and standard Python documentation, all of which contribute to enhanced maintainability and reproducibility.

#### 2.1.3 SPAC Interactive Analysis Layer

The SPAC ecosystem can be deployed on any enterprise platform, such as Code Ocean [26] or Code Workbook from Palantir Foundry [27]. These platforms provide robust authentication and a configurable computational graph to manage workflows efficiently. For our deployment, we selected Code Workbook due to its modular building blocks that execute code, generate visualizations, output datasets, and record logs. For intramural NIH research scientists and their collaborators, the NIH Integrated Data Analysis Platform (NIDAP) [28] built on Palantir Foundry has been tailored for non-proprietary code and workflow development. By leveraging NIDAP’s Code Workbook templates and pipelines, the SPAC Interactive Analysis Layer exposes the underlying SPAC Python Package and its dependencies in a user-friendly, web-based environment. Within this framework, analysts can construct stepwise pipelines to ingest data, apply preprocessing or quality control, run clustering or spatial analysis, and generate publishable figures. Key features include:

- **Web-Hosted Pipelines:** Configure parameters, run workflows, and monitor progress entirely through a browser interface, eliminating the need for local installations.
- **HPC Integration and Scalability Enhancements:** SPAC seamlessly scales analyses to HPC environments with automated workload distribution for large datasets. In our implementation, we deployed SPAC on Biowulf [29], the on-premise NIH HPC cluster, which supports distributed or parallel execution (e.g., on GPU-enabled nodes). This ensures that computationally intensive steps such as iterative clustering or large-scale image data processing, can be performed efficiently.
- **Collaboration and Reporting:** Compile final results into shareable, “technical-free” reports that detail each analysis step and associated figures, promoting transparency and reproducibility across research teams.

#### 2.1.4 SPAC Real-Time Visualization Layer

At the top of the stack, the SPAC Real-Time Visualization Layer leverages Shiny Application for python [30] with Posit Connect [31]to deliver on-the-fly interactive dashboards and exploratory figures. By decoupling heavy computations from the visualization front end, this layer empowers principal investigators, bench scientists, and other non-specialist stakeholders to quickly explore large spatial datasets—toggling between features or annotations and drilling down into specific regions of interest—while simultaneously generating live dashboards that create dynamic views, such as UMAP plots, hierarchical heatmaps, and neighborhood graphs, which update in real time as new data are loaded or parameters are adjusted.

This four-tier architecture establishes a clear separation of interests between development and sharing. Software engineers and data scientists can concentrate on code quality and methodological innovation, while biologists and clinicians benefit from an intuitive, browser-based environment for data exploration and interpretation. In practice, image analysts or bioinformaticians design custom analyses using the SPAC Python Package; core facility scientists or postdoctoral researchers run complex pipelines via the Interactive Analysis Layer; and principal investigators or bench scientists rely on the Real-Time Visualization Layer to review outcomes. By integrating open-source solutions with enterprise-ready HPC scaling, SPAC effectively streamlines the workflow from raw data to actionable insights in single-cell spatial proteomics research.

### 2.2 Modules and NIDAP template

Fig. 3 provides an overview of the SPAC workflow for single-cell spatial analysis, while Supplementary Fig.1 presents a representative NIDAP Code Workbook that demonstrates the typical SPAC workflow. This workflow encompasses data aggregation, multiple-slide data sampling, exploratory data analysis, feature preprocessing and normalization, clustering, dimensionality reduction, cluster analysis, and spatial analysis. We elaborate on four key modules that collectively streamline these steps into an efficient and reproducible analysis pipeline.

**Fig. 3.**
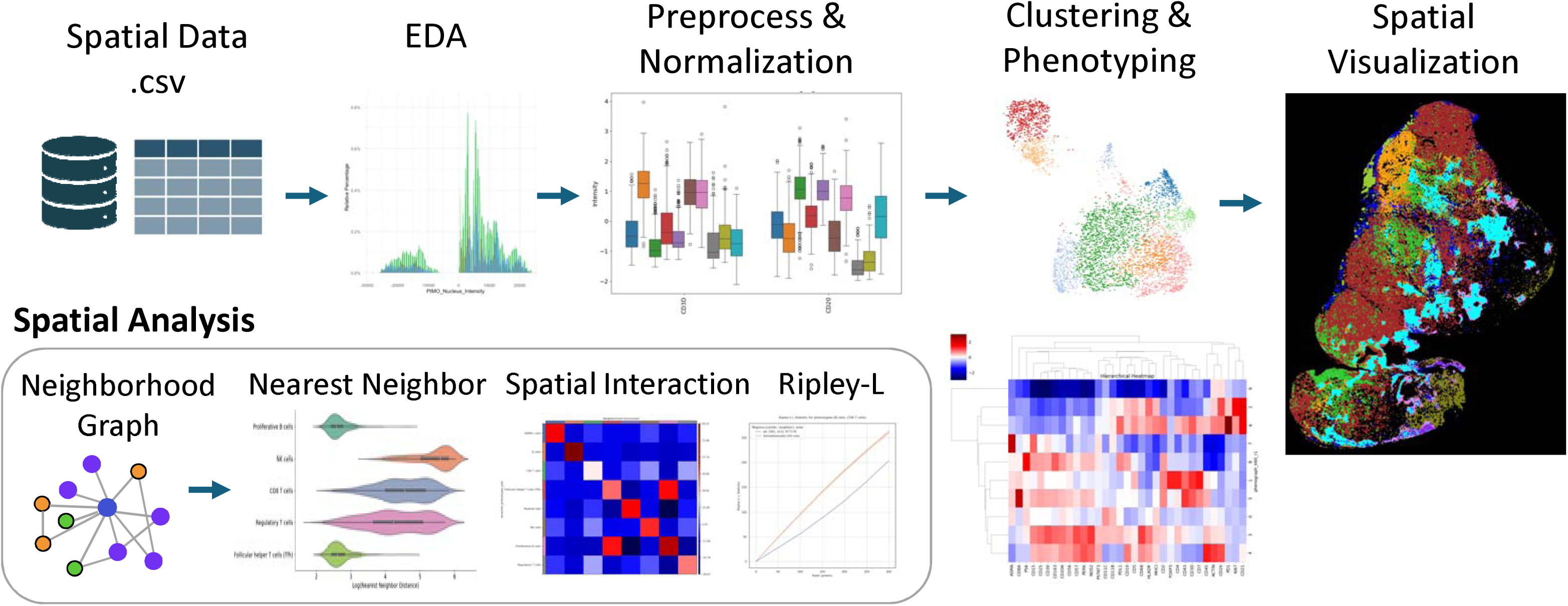
An overview of the SPAC Workflow for single-cell spatial analysis. The top row depicts the main pipeline steps: In the **Spatial Data** step, CSV files are imported, merged, and optionally downsampled or annotated. **Exploratory Data Analysis (EDA)** uses methods such as histograms and boxplots to reveal marker distributions and potential outliers. During **Preprocessing & Normalization**, techniques like quantile scaling, arcsinh transformation, and batch corrections are applied to standardize the data. **Clustering & Phenotyping** relies on algorithms like PhenoGraph, t-SNE, or UMAP for grouping cells by marker expression and assigning phenotypes. In **Spatial Visualization**, these phenotypes are color-coded and overlaid on tissue images. The bottom row focuses on **Spatial Analysis** modules, including building a Neighborhood Graph to depict relationships among cells, performing Nearest Neighbor analyses to quantify infiltration or proximity, detecting Spatial Interactions between cell types, and calculating Ripley’s L to measure clustering or dispersion over distance scales. A **Hierarchical Heatmap** (at right) integrates marker expression or phenotype relationships, providing deeper insights into subpopulations and their spatial patterns.

#### 2.2.1 Key Data Structure and Formats

In multiplexed tissue imaging workflow, the initial data are generated by image-processing software (e.g., HALO, MCMICRO, Visiopharm [32], QuPath [33]) that performs cell segmentation and computes per-cell features such as protein staining intensities, morphological descriptors (e.g., size, eccentricity), and spatial coordinates. These measurements are typically exported in tabular format (often CSV files), where each row corresponds to a single cell, and columns capture intensities, metadata, and positional attributes.

Users can batch-upload these files into the NIDAP platform, which aggregates multiple files into a single dataset while enforcing granular access control at both group and individual levels. Upon ingestion, SPAC converts these single-cell tables (and any optional annotations) into AnnData objects—a widely adopted data structure in single-cell genomics and proteomics. This standardized format facilitates the storage of multiple data versions by allowing raw, normalized, and annotated layers to coexist within a single object, and it enables seamless interoperability by integrating directly with the broader single-cell Python ecosystem (e.g., SCANPY, Squidpy), thereby streamlining downstream analysis and visualization without cumbersome file transformations. Additionally, by retaining X–Y positional metadata, the format readily supports spatial operations such as neighborhood graph construction, distance computations, and interactive map overlays, ultimately ensuring consistency throughout preprocessing, clustering, and spatial analysis steps while simplifying collaboration and transitioning between exploratory and production workflows.

#### 2.2.2 Visualization of Features and Annotations

In SPAC, two fundamental terms are used to describe the data: Features and Annotations. Features are measurable attributes of each cell, such as biomarker intensity. Annotations are categories assigned to cells to group them based on specific criteria. For example, annotations can denote cell type (e.g., “animal type” with labels like “animal1” and “animal2”), spatial regions (e.g., “hypoxic”, “normal stroma”), or clusters derived from computational methods (e.g., “phenograph”), thereby facilitating both biological interpretation and spatial analysis.

SPAC provides a comprehensive suite of integrated visualization tools that span the entire workflow from exploratory data analysis to final result presentation. Implemented using libraries such as Matplotlib, Plotly, and Seaborn, SPAC provides basic plots (e.g., histograms, boxplots, heatmaps, and scatter plots) to examine features and annotations, revealing underlying patterns, distributions and relationships. Built upon these standard charts, SPAC enables visualizations tailored for high-dimensional data, spatial statistics and spatial organization. For example, dimensional reduction plots project complex, multi-feature data into lower-dimensional representations, thereby making it easier to discern overall similarities and differences among cells or samples. Hierarchical heatmaps with dendrogram, on the other hand, visualize hierarchical clustering of feature or annotation dimension based on computed correlation matrix, elucidating the hierarchical structure of cellular subpopulations. These visualization modules are embedded directly into the analysis pipeline, ensuring that results at each stage can be immediately inspected and interactively explored without the need to export data to external software.

#### 2.2.3 Cell Phenotypes, States and Functions

A central objective in single-cell spatial proteomics is to identify and characterize diverse cell phenotypes. Multiplexed imaging platforms capable of measuring 10 to 60 markers per cell, are well suited for dissecting heterogeneous cell populations from broad lineages (e.g., immune vs. epithelial) to subtle functional states (e.g., signaling pathway activation or T-cell exhaustion). SPAC supports two complementary strategies for phenotypes determination: knowledge-based (manual gating) and data-driven (unsupervised clustering) approaches.

The knowledge-based phenotyping approach leverages well-established biomarkers (e.g., immunological or histopathological markers) associated with specific cell types and manually defines phenotypes by setting intensity-based thresholds. Marker expressions are categorized into binary states (e.g., ‘0’ or ‘1’) to represent phenotypic classes such as “CD4-negative” versus “CD4-positive,” allowing precise, context-specific annotation of cell populations. SPAC can also consolidate multiple biomarker expressions into a single composite phenotype_code. For example, a regulatory T-cell phenotype can be succinctly defined as CD4+CD25+FOXP3+, integrating simultaneous expressions of multiple markers into one standardized code. This flexibility enables domain experts to annotate specific cell subtypes or functional states without custom programming, simply by specifying a human-readable phenotype_name and the corresponding composite phenotype_code. By providing standardized and reproducible phenotype definitions, this feature enhances scalability and ensures consistency across extensive, multi-slide datasets, which is particularly valuable in high-throughput experimental contexts.

Complementing this manual approach, SPAC supports unsupervised clustering algorithms, such as PhenoGraph, to systematically uncover novel cell populations based purely on phenotypic similarity. PhenoGraph employs a three-step process: it first normalizes the marker intensity data to ensure comparability across cells, then constructs a k-nearest neighbor (KNN) graph by identifying the k nearest neighbors for each cell using Euclidean distance, and subsequently refines the graph by adjusting the edge weights such that the weight between any two cells scales with the number of neighbors they share. Finally, graph clustering using either the Louvain or Leiden algorithm is applied to delineate distinct cell communities. With adjustable clustering resolutions, this method can reveal subtle subpopulations often missed by manual gating. Additionally, dimensionality reduction techniques (e.g., t-SNE or UMAP) provide visual summaries of high-dimensional data, facilitating inspection of emergent clusters and guiding hypothesis generation. These unsupervised methods can identify phenotypic clusters associated with biological or clinical outcomes such as patient prognosis, thus enabling refined patient stratification beyond conventional clinical classifications.

By integrating manual gating methods anchored in established biomarker knowledge with flexible unsupervised clustering techniques, SPAC provides a robust and versatile framework that examines known cell phenotypes under various states or conditions, and simultaneously discovers novel subpopulations, advancing our understanding of tissue organization and disease mechanisms.

#### 2.2.4 Spatial Analysis

SPAC streamlines the analysis of spatial organization within multiplexed tissue images across multiple scales, from nearest neighbor distances to larger scale clustering patterns. It integrates spatial analytic methods with intuitive visualization tools, and systematically interrogates spatial relationships among cell phenotypes, such as assessing phenotype arrangements, identifying co-location or exclusion patterns, and detecting significant cell–cell interactions beyond random chance.

**Nearest Neighbor analysis** quantifies the proximity of a specified “source” phenotype relative to other cell types, highlighting patterns of local adjacency or avoidance. To assess broader spatial patterns, particularly when comparing Case versus Control conditions, SPAC implements **Ripley’s L statistic** [34] measuring whether two phenotypes exhibit spatial aggregation, dispersion, or random distribution across varying spatial scales, where the random distribution is modeled using a Poisson point process under the assumptions of homogeneous tissue distribution. SPAC builds **Neighborhood Graphs** using Squidpy by connecting cells via *k*-nearest neighbors or radius-based criteria, computes **Cluster Interaction Matrix** (tallying the edges between distinct phenotypes), and applies permutation-based **Neighborhood Enrichment** scores to determine whether phenotype pairs co-locate significantly more or less frequently than expected by chance.

In addition, SPAC includes a customizable neighborhood profiling framework that captures the local cellular environment around each cell. This feature summarizes the composition of a cell’s immediate neighborhood, providing a high-resolution context for every single cell. Such neighborhood profiles enable flexible and scalable downstream analyses, for example generating a spatial UMAP embedding [35] that groups cells by similarity of their microenvironments, or performing infiltration profiling to quantify how certain cell types penetrate into different tissue regions. By characterizing each cell’s neighborhood across different samples or conditions, researchers can interpret spatial organization differences with unprecedented resolution, gaining deeper insight into tissue structure and cell–cell interactions.

Spatial plotting capabilities—both static and interactive—enable visualization and refinement of spatial distribution within defined tissue contexts. The interactive spatial plot, built with Plotly Express, supports real-time exploration features, such as zooming, on-hover annotations, and pined-color mapping for consistent and customizable phenotype labelling, facilitating clear interpretation and shareable visualizations. Together, these analytic and visualization tools form a comprehensive suite for dissecting complex spatial topographies within multiplexed tissue samples.

### 2.3 HPC Connector for Scaling

A core objective of SPAC is to address large-scale computational demands by seamlessly offloading resource-intensive analyses to HPC clusters (Fig. 4). The NIDAP HPC Connector integrates HPC resources directly into the interactive analysis layer, allowing any user-configured computational step—such as PhenoGraph clustering or other advanced analyses— to be executed on an HPC node (CPU or GPU). When a computationally demanding task is initiated, the connector automatically identifies the corresponding input data within NIDAP, dispatches the job to the appropriate HPC resource, and monitors job status in real time through an integrated HPC Dashboard. Upon completion, results are seamlessly returned to the interactive interface, enabling rapid, large-scale analyses without requiring specialized HPC credentials or command-line interaction.

**Fig 4.**
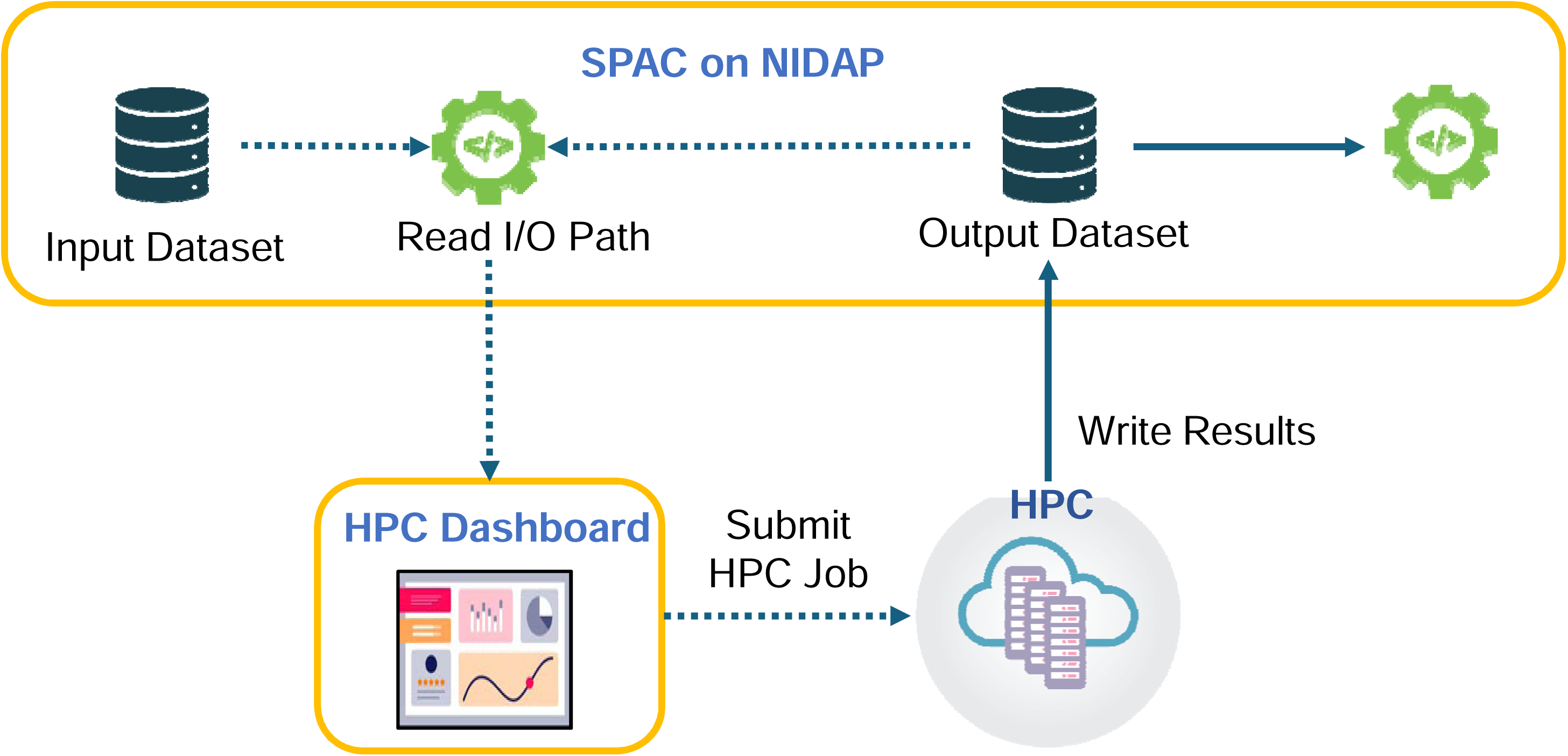
Scaling Single-Cell Analysis on HPC Clusters via the NIDAP HPC Connector. SPAC leverages the NIDAP HPC Connector to offload large single-cell analyses to HPC clusters. A secure service account reads input data from the SPAC pipeline on NIDAP, then submits the job to GPU or high-memory CPU nodes via HPC Dashboard. The HPC Dashboard provides real-time monitoring of job status, resource utilization, and logs. Once computations complete, results are automatically returned to NIDAP for further visualization and downstream analysis. This workflow enables large-scale computational tasks without requiring users to have direct HPC credentials or command-line interaction.

#### 2.3.1 Infrastructure for Multithreading and Parallel Processing

Under the hood, the HPC Connector leverages the high computational resources available on institutional clusters (e.g., Slurm on NIH Biowulf) to distribute large workloads across multiple nodes or GPUs. For example, a PhenoGraph clustering job that might take an hour (or potentially fail) on a desktop environment can be completed in under ten minutes using GPU nodes. This approach not only scales to datasets comprising millions of cells but also provides a robust foundation for future integration of advanced machine learning and algorithmic tasks.

#### 2.3.2 Usability

From the user’s perspective, HPC scaling is simplified to selecting a desired compute mode— CPU or GPU—within the Code Workbook interface. The HPC Code Workbook Connector processes the request and submits it to the appropriate HPC partition (e.g., “quick,” “norm,” or “GPU”) with the necessary resources (e.g., core count, memory, time). Logging, error checking, and intermediate file handling are automatically relayed back to the user’s Code Workbook node, ensuring a streamlined experience without direct HPC account management.

#### 2.3.3 Speed

By leveraging powerful cluster nodes with up to terabytes memory [29] capacities and extensive GPU arrays, the HPC Connector significantly reduces execution times for large SPAC jobs. Bottlenecks typically associated with single-threaded operations, such as clustering, dimensionality reduction, or spatial interaction statistics, are mitigated through effective parallelization. Benchmarks on synthetic datasets (up to five million cells with approximately 20 features each) demonstrate near-linear scalability when running PhenoGraph at k=30 (number of nearest neighbors) and resolution=1.0 (cluster granularity), as shown in Fig. 5. This setup enables users to explore multiple clustering parameters within a single session while maintaining rapid execution times.

**Fig 5.**
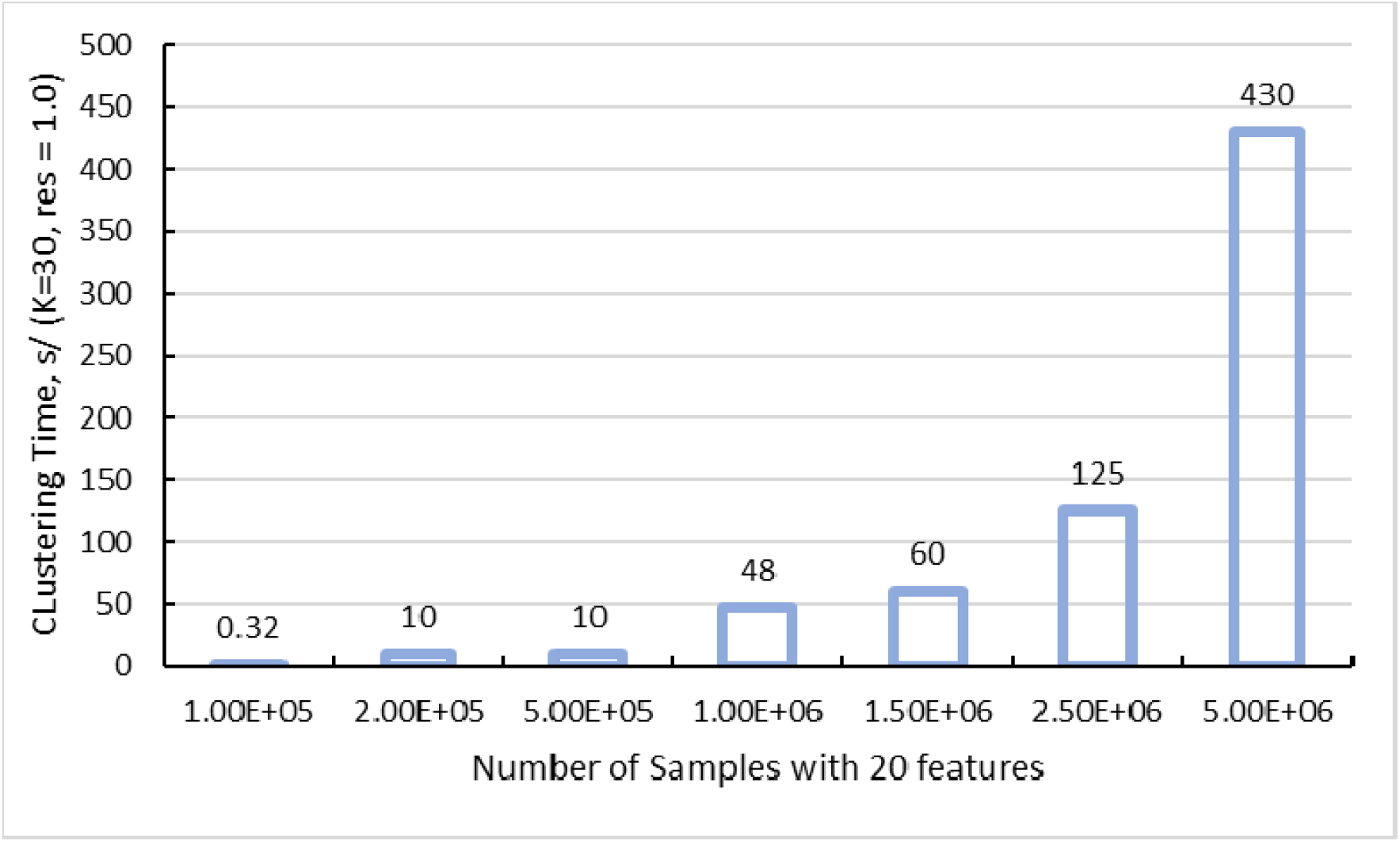
GPU-Accelerated PhenoGraph Clustering Times on Synthetic Datasets (k=30, resolution=1.0). The bar chart illustrates how clustering runtime scales when applying GPU-accelerated PhenoGraph to synthetic single-cell datasets ranging from 100,000 up to 5 million cells, each with 20 features. The parameter k=30 specifies the number of nearest neighbors used to construct the graph, while resolution=1.0 determines the granularity of cluster detection. Even at the upper limit of five million cells, total processing time remains on the order of minutes, demonstrating near-linear scalability and underscoring the efficiency gains from leveraging HPC resources.

#### 2.3.4 Security

The HPC Connector employs a “service account” framework to ensure data integrity and security. Instead of requiring personal HPC credentials, the NIDAP platform securely manages job submission and file transfers via established, audited APIs. All data transfers occur within a controlled environment using encrypted connections, thereby preventing unauthorized access to sensitive research data. This approach upholds enterprise-level security while providing the scalable compute infrastructure essential for spatial single-cell analysis. It is important to note that this environment is not authorized for use with personally identifiable data.

## 3. Results

To evaluate SPAC’s performance on real-world data, we applied the SPAC across multiple collaborative research projects, using hundreds of slides and tens of millions of cells. Here, we highlight one project investigating the impact of hypoxia on cellular neighborhoods within the tumor microenvironment in a syngeneic mouse model of triple negative breast cancer using 4T1 cells [36]. To delineate hypoxic regions, pimonidazole (PIMO) was administered via tail vein injection at a dose of 60 mg/kg, 30 minutes prior to euthanization and tissue harvest. An end-to-end analysis was performed using the workbook of SPAC Interactive Analysis Layer along with the SPAC Interactive Dashboard, a Python-based Shiny application for real-time visualization and sharing of results. This showcase demonstrates SPAC’s versatility in characterizing cell phenotypes and analyzing spatial distribution in multiplex imaging data, with additional spatial interaction, biological interpretation and statistical analyses to be presented elsewhere.

The single-cell dataset was derived from MxIF images processed using HALO and comprised approximately 2.6 million cells. Each cell was characterized by its spatial coordinates, intensities for nine biomarkers (Hif1a, NOS2, COX2, β-catenin, vimentin, E-cadherin, Ki67, PIMO, aSMA), morphological features (e.g., cytoplasm area, nucleus roundness), and localization within three distinct tumor regions: tumor normoxia (median oxygen level: 6.8%), tumor hypoxia (median oxygen level: 1.3%) [37] and necrotic areas. Whole tissue slides were collected from eight animals at three time points (days 9, 19, 30 after tumor cell injection).

### 3.1 GPU Acceleration of Unsupervised Clustering and Optimization

Building on the computational challenges outlined earlier, GPU resources provide advantages for tasks dominated by matrix operations and graph-based optimizations (e.g. PhenoGraph clustering, Louvain clustering, and UMAP dimensionality reduction). In this study, an unsupervised PhenoGraph clustering task on 2.6 million cells with nine biomarkers was accelerated from approximately 2.5 hours on an AMD Epyc 7543-based CPU node to about 7 minutes on a NVIDIA A100 GPU node using the Grapheno implementation [38]. The CPU run utilized an average of 7.5 cores with 13 GB of memory. This resulted in a ∼20-fold speedup, highlighting the efficiency of GPU acceleration for large-scale clustering tasks.

A key factor in the GPU’s performance is its streamlined workflow. The implementation begins with a single k-nearest neighbors (KNN) calculation that construct a connectivity graph by efficiently capturing local similarities in high-dimensional cell data. This graph then serves as the basis for four separate Louvain clustering procedures executed at different resolution parameters. Louvain clustering is selected for its robustness in detecting community structures and its adaptability in revealing clusters at varying granularities. Because of inherent parallelization and efficient GPU memory management, running multiple clustering resolutions adds only minimal overhead compared to computing each independently. Thus, the entire process—from the KNN calculation through all clustering steps—is completed in just a few minutes, highlighting the substantial acceleration achieved over traditional CPU-based methods.

To optimize cluster granularity, we systematically varied the PhenoGraph parameters: the number of neighbors (k) and the resolution, which controls the coarseness of clustering. As illustrated in Fig.6, SPAC’s HPC pipeline enabled rapid exploration of these parameter settings.

**Fig 6.**
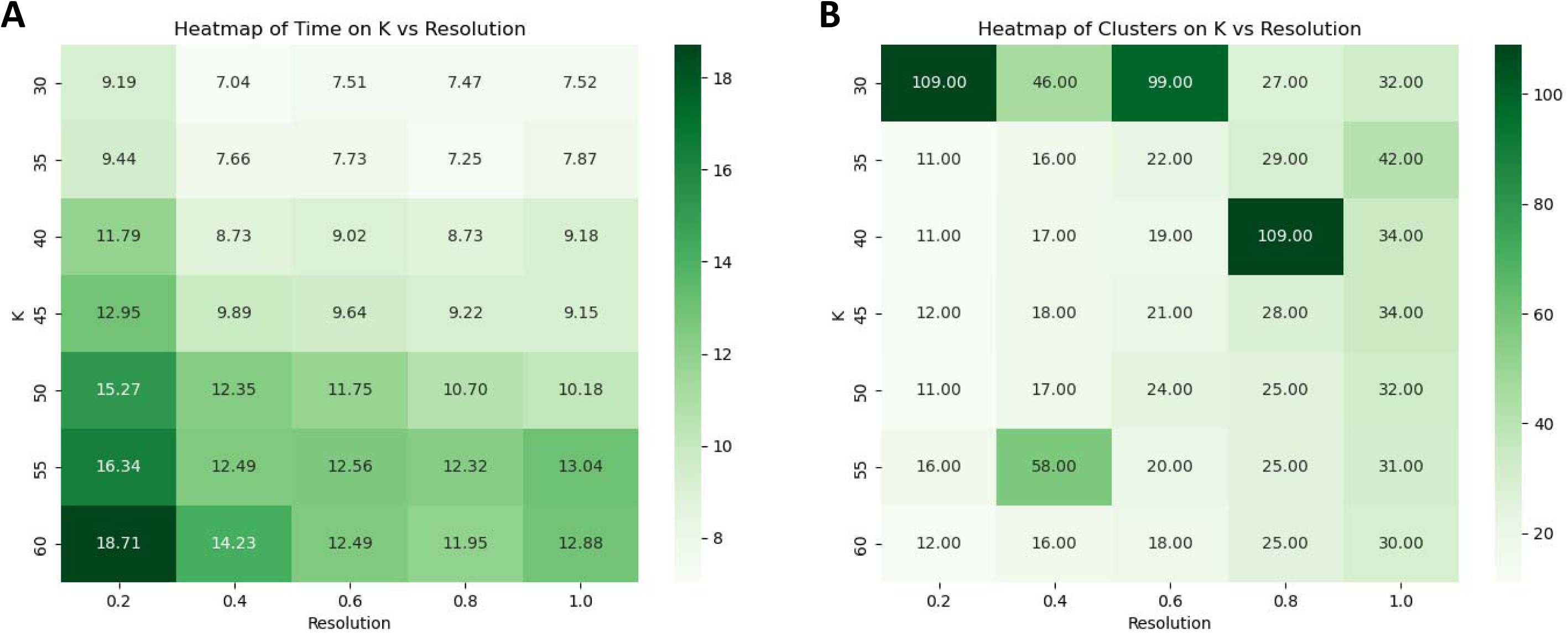
GPU-Accelerated PhenoGraph Tuning Across k-neighbors and Resolution. (**A)** Heatmap of runtime (minutes) across combinations of PhenoGraph parameters k (vertical axis) and resolution (horizontal axis), using ∼2.6 million cells with 9 biomarkers. Darker green cells indicate longer runtimes, although all configurations remain under 20 minutes. (**B)** Corresponding heatmap of the total number of clusters produced. In this parameterization, a larger numeric resolution value yields a more granular partition, producing more clusters. These results highlight SPAC’s ability to leverage HPC GPU resources and systematically explore parameter space with interactive turnaround times, enabling fine-grained analysis of tumor-associated cell populations.

The runtime heatmap (Fig. 6A) shows that all combinations of k and resolution completed in under 30 minutes on a GPU, while the cluster-count heatmap (Fig. 6B) reveals that higher resolution values yield a greater number of clusters. By leveraging GPU acceleration and parallel computing, SPAC allows users to test multiple configurations within a single session.

### 3.2 Phenotype Annotation

The outputs from PhenoGraph clustering across various parameter settings were evaluated using hierarchical heatmap of biomarker expression and UMAP visualization, alongside expert domain knowledge. The optimal condition (k=35, resolution=0.6) yielded 16 clusters, a choice driven by the balance between granularity and interpretability. Specifically, this setting effectively captured distinct cellular subpopulations while minimizing noise and over-segmentation. Each of the 16 clusters exhibited a unique biomarker expression profile, which was consistent with established cellular phenotypes, thereby facilitating annotation and renaming for downstream analysis.

For example, the hierarchical heatmap (Supplementary Fig. 2), displaying z-score normalized marker expressions, revealed that clusters 4 and 15 with magenta boxes due to their uniquely high levels of E-cadherin and β-catenin. These clusters were consequently merged and annotated as “Ecad+b-catenin+”. Similarly, clusters 0 and 7, marked by red boxes, exhibited moderate expression of PIMO, a marker indicative of hypoxia; these were combined and renamed “Hypoxic_PIMO-dim”. The final annotated phenotypes are presented in the hierarchical heatmap in Fig. 7, where the merged phenotypes are again outlined in the same color scheme to facilitate direct comparison.

**Fig. 7.**
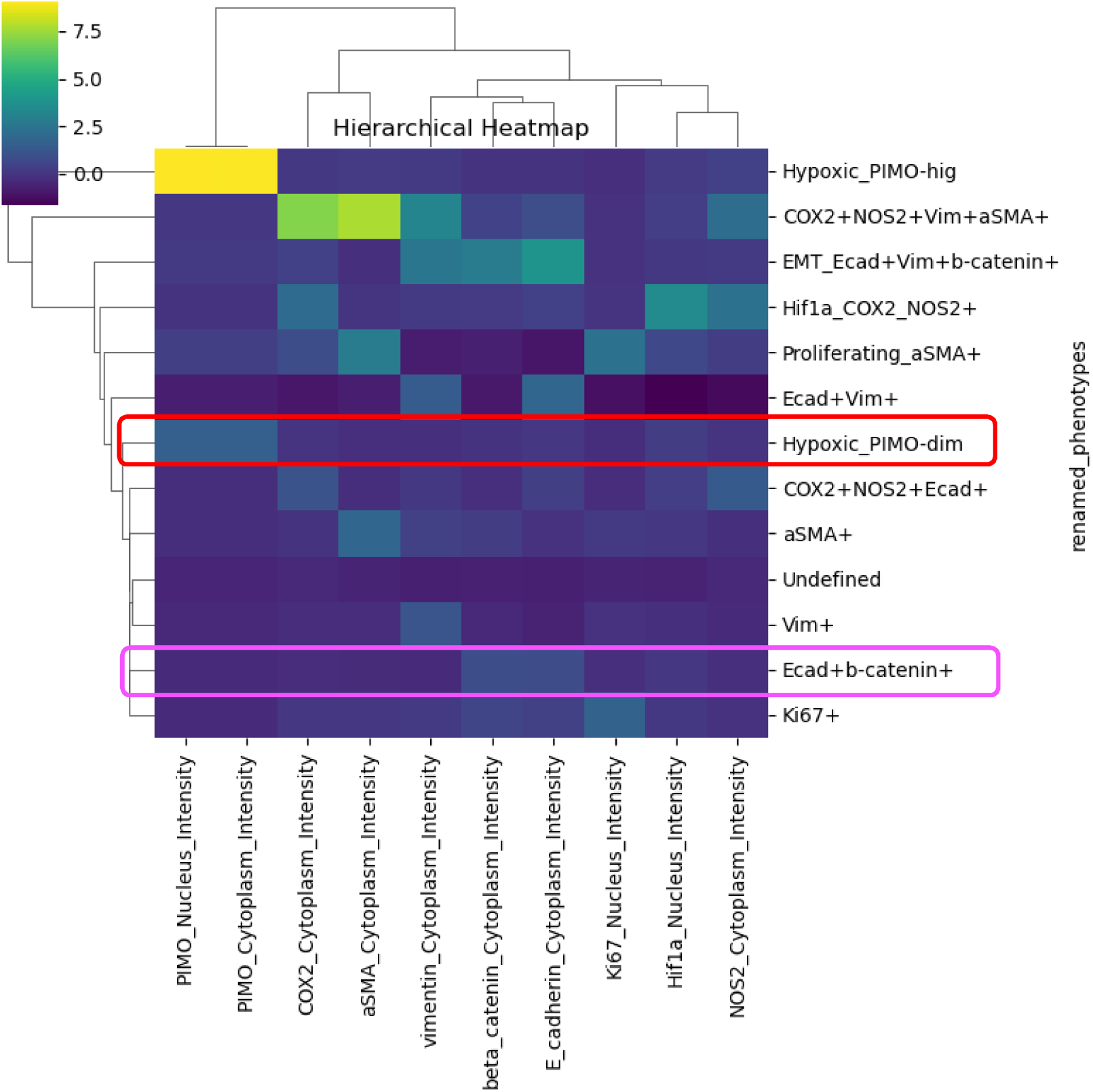
Hierarchical heatmap of renamed phenotypes (rows) clustered by marker intensities (columns). Each cell represents the scaled expression level (blue=lower expression; yellow=higher expression) of a particular marker within a given renamed phenotype. Dendrograms indicate similarity groupings among both markers and phenotypes, revealing distinct subpopulations based on their shared expression patterns. Clusters 4 and 15 (outlined in magenta in Supplementary Fig. 2) are merged into the “Ecad+b-catenin+” phenotype. Clusters 0 and 7 (outlined in red in Supplementary Fig. 2) form the “Hypoxic_PIMO-dim” phenotype.

### 3.3 Spatial Distribution of Cellular Phenotypes Across Tumor Regions

To further illustrate SPAC’s capabilities, we leveraged the SPAC Interactive Dashboard to investigate the spatial distribution of cellular phenotypes across tumor regions. We hypothesized that epithelial cells characterized by high expression of E-cadherin and β-catenin are predominantly localized within tumor normoxic regions, reflecting the epithelial nature of 4T1-induced tumors. In contrast, cells annotated as “Hypoxic_PIMO-dim” were expected to reside primarily in tumor hypoxic regions.

Two complementary visualization tools were deployed. First, the Relational Heatmap employs color gradients to depict the frequency distribution between categorical annotations (i.e., tumor region and cellular phenotype). For instance, the heatmap (Fig. 8, top) indicates that 53.2% of cells in tumor hypoxic regions are classified as Hypoxic_PIMO-dim, whereas 16.1% of cells in tumor normoxic regions express E-cadherin/β-catenin. Second, the Sankey Plot provides a flow diagram illustrating the proportional connectivity between tumor regions and phenotype annotations. In this display (Fig. 8, bottom), between 53.2% and 62.6% of Hypoxic_PIMO-dim cells originate from tumor hypoxic regions (representing approximately 312,000 cells), thereby reinforcing the expected spatial segregation. All analyses presented here were performed on a combined dataset comprising all animals.

**Fig. 8.**
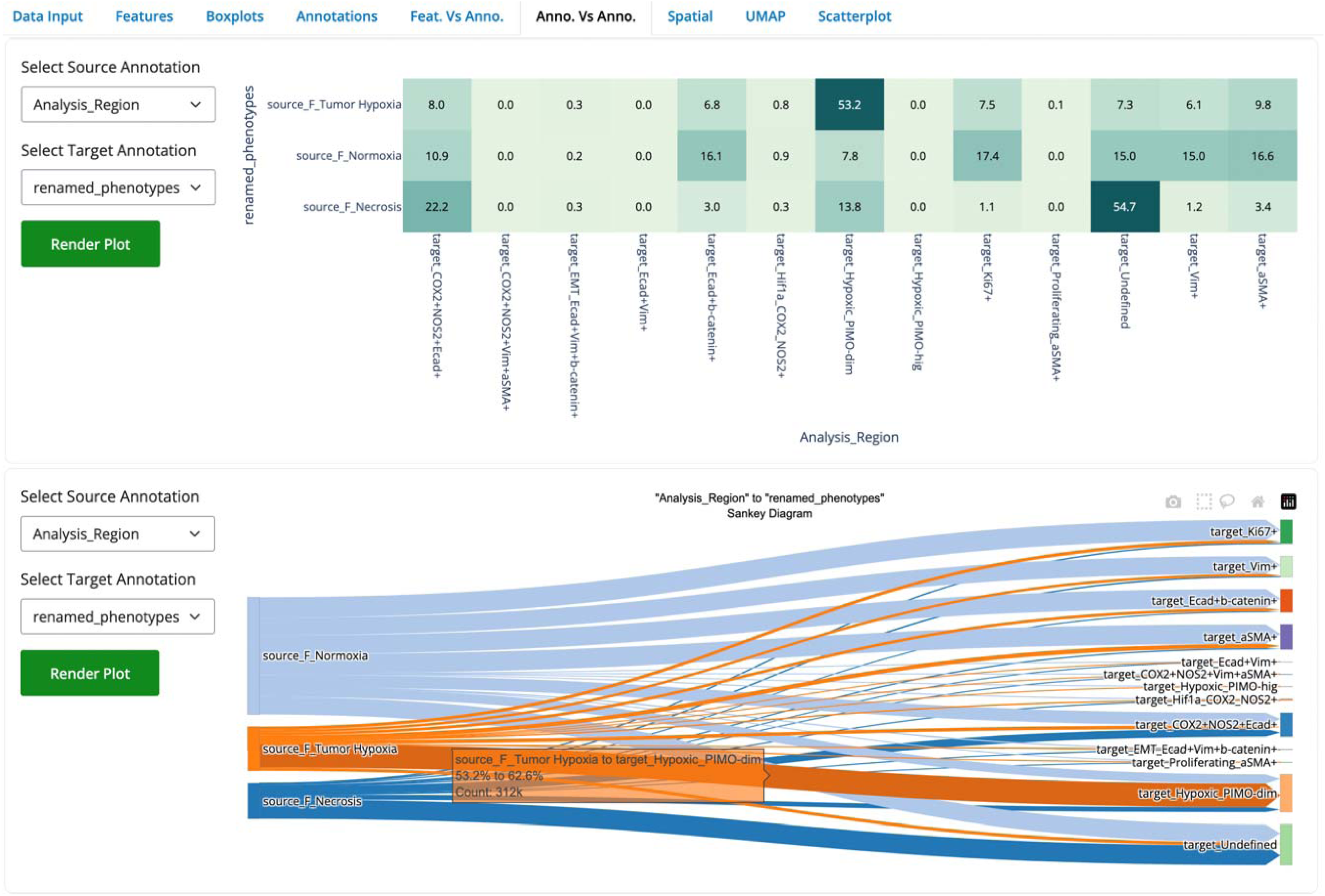
Relational Heatmap and Sankey Plot depicting the distribution of cells from different tumor regions (source annotation) across renamed cellular phenotypes (target annotation). In the Relational Heatmap (top), each cell of the matrix displays the percentage of cells (row total = 100%) that share both a specific tumor region (e.g., Hypoxia, Normoxia, Necrosis) and a given phenotype (e.g., Hypoxic_PIMO-dim, E-cadherin/β-catenin). Notably, 53.2% of cells in tumor hypoxic regions are Hypoxic_PIMO-dim, while E-cadherin/β-catenin– positive cells comprise approximately 16.1% of the normoxic compartment. The Sankey Plot (bottom) visualizes these same relationships as proportional flows, indicating, for example, that 53.2–62.6% of all Hypoxic_PIMO-dim cells originate from hypoxic regions (∼312,000 cells). Both the heatmap and Sankey diagram are generated by SPAC’s interactive dashboard, enabling dynamic exploration of cellular annotation and spatial context.

SPAC’s Spatial Plot tools enable the projection of cellular phenotypes onto the original spatial coordinates of tissue sections, offering both static and interactive modes for visualizing single-cell distributions. The Interactive Spatial Plot provides dynamic user-driven exploration of cell populations across multiple annotations, facilitating deeper biological interpretation in real time. As illustrated in Fig. 9, three distinct tumor regions, normoxic, hypoxic, and necrotic are defined in panel-A. Panel-B highlights epithelial cells co-expressing E-cadherin and β-catenin, which localize predominantly within normoxic areas, while panel-C shows “Hypoxic_PIMO-dim” cells enriched in hypoxic and necrotic regions. In panel-D, the bar chart quantifies these distributions at day 30, revealing that “Hypoxic_PIMO-dim” cells are predominant in hypoxic regions, whereas the epithelial population remains largely confined to normoxic regions.

**Fig. 9.**
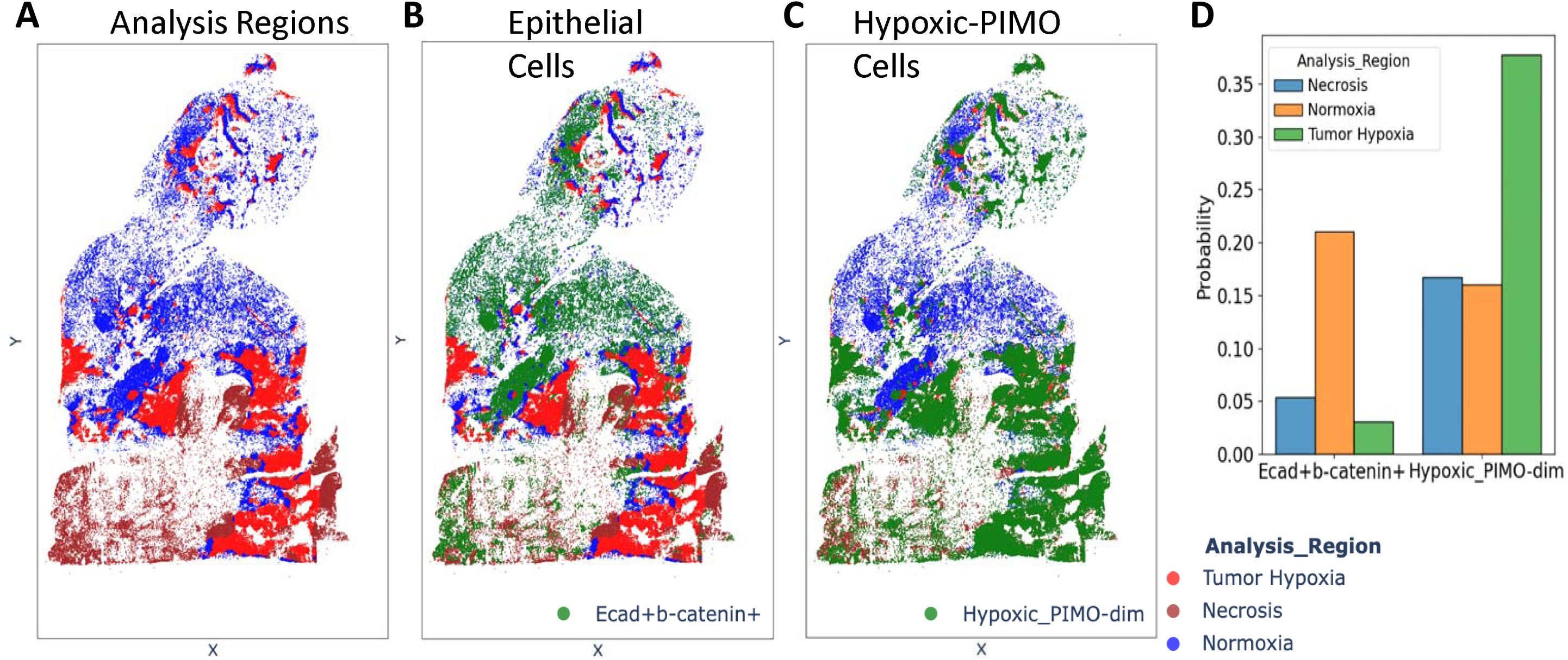
Spatial distribution of E-cadherin/β-catenin expressing epithelial cells and hypoxic-PIMO cells across different tumor regions at day 30. **(A)** Three tumor regions (normoxia, hypoxia, and necrosis) delineated for spatial analysis. **(B)** E-cadherin+β-catenin+ epithelial cells (green) predominantly localize in the normoxic region (blue), indicating higher oxygen levels. **(C)** Hypoxic_PIMO-dim cells (green) are enriched in the hypoxic (green) and necrotic (red) regions, reflecting reduced oxygen availability. **(D)** Bar chart summarizing the proportion of E-cadherin+β-catenin+ and Hypoxic_PIMO-dim cells across the three tumor regions. The results highlight distinct spatial patterns for epithelial and hypoxic cell populations at day 30.

This overlay analysis underscores SPAC’s ability to delineate tumor subregions based on distinct cellular phenotypes, an observation consistent with the underlying biology of the 4T1 model, where epithelial markers typically localize to normoxic areas, whereas hypoxic regions will preferentially harbor hypoxia markers associated with PIMO. Notably, SPAC supports the simultaneous display of multiple annotations (e.g., region labels and cell phenotypes) within each spatial plot, along with user-defined pin colors and stratification options. Such capabilities remain challenging in many existing tools but are essential for capturing the complexity of tumor microenvironments and facilitating intuitive, high-throughput analysis.

Collectively, these findings demonstrate that SPAC robustly integrates phenotype annotation with high-resolution spatial visualization, providing a powerful tool for deciphering the complex spatial organization of tumor tissues. This integrated approach not only streamlines analysis but also lays the groundwork for deeper insights into tumor biology and microenvironmental interactions.

## 4. Discussion

SPAC was developed to meet the growing need for an integrated, user-friendly platform that streamlines the analysis of high-dimensional, spatially resolved single-cell data from multiplex imaging. By combining a modular software architecture, a web-based interactive interface, and robust HPC integration, SPAC supports comprehensive end-to-end analysis through intuitive graphical workflows that are accessible to both computational experts and experimentalists. Its interactive dashboards provide real-time visualization of plots and parameter adjustments, enabling immediate feedback and facilitating data sharing.

In a collaborative study with research groups at the National Cancer Institute’s Frederick National Laboratory (NCI/FNL), SPAC demonstrated its practical utility. For example, in a 4T1 breast cancer model, SPAC processed a dataset of 2.6 million cells and reduced unsupervised clustering runtime from eight hours on a CPU to under ten minutes on a GPU. Its ability to rapidly optimize clustering, phenotyping, and visualize spatial distributions allowed clear delineation of tumor regions, for instance, distinguishing normoxic areas enriched in E-cadherin/β-catenin-positive cells from hypoxic regions dominated by hypoxia-associated cells, thereby providing data-driven insights into tissue organization and tumor microenvironment heterogeneity.

To date, our collaborations with eight research groups at NCI/FNL, involving the processing of over 30 million cells across diverse studies, underscore SPAC’s ability to bridge the gap between bench scientists and data analysts. Through collaboration with the Purdue Institute for Cancer Research, the SPAC Python library was integrated into a Shiny application layer, and contributions from the Purdue Data Mine (https://datamine.purdue.edu) further expanded its functionality. Collectively, these efforts highlight SPAC’s growing adoption and usability in a variety of research settings.

Looking forward, we are actively developing several enhancements. Planned improvements include the integration of spatial transcriptomics analysis for the dataset from technologies such as Visium [39] and Slide-Seq [40], expanding quality control and statistical hypothesis testing for case-control comparisons, and incorporating additional clustering algorithms along with further GPU acceleration. We also aim to extend SPAC’s spatial analysis capabilities by adding advanced spatial interaction metrics and refined batch correction strategies. One particularly exciting avenue is the adoption of the Vitesse framework, which integrates downstream analysis with high-resolution multiplex slide images into a unified, web-accessible dashboard, thereby further streamlining data interpretation and sharing.

## 5. Conclusion

In conclusion, SPAC addresses key challenges in spatial single-cell proteomics by providing an efficient, scalable, and user-friendly platform for high-throughput analyses of multiplex imaging data. Its integration of HPC acceleration with interactive visualizations enables rapid parameter optimization and detailed spatial characterization of cellular phenotypes. This capability is valuable for clinical applications, where comparative analyses across primary tumors, metastases, or samples with distinct genomic profiles, and their relationships to immune infiltration and therapy response, are critical. Ultimately, SPAC not only enhances our understanding of tissue organization and its clinical implications but also fosters a more integrated and collaborative approach to spatial single-cell analysis.

## Abbreviations

SPAC: Analysis of SPAtial Single-Cell
NIDAP: NIH Integrated Data Analysis Platform
HPC: High-Performance Computing
WSI: Whole-Slide Tissue Imaging
MxIF: Multiplexed ImmunoFluorescence

## Acknowledgements

We would like to thank Tyler Malys for suggestions on batch normalization, Alex Mitrophanov for exploring Ripley-L method, and thank our collaborators and members of the NCI/FNL, including Noemi Kedei, Scott Lawrence, Thomas Hu, Milind Pore, Paul Mallory, Will Heinz, Andrew Weisman, Andrei Bombin, Wiem Lassoued, Mike Kelly, and Christof Kaltenmeier for their invaluable feedback.

Thanks to the single-cell and spatial imaging communities for inspiring a tool that bridges the gap from datasets to knowledge, accelerating discoveries and innovation.

Palantir Foundry was used in the integration, harmonization, and analysis of SPAC on the hypoxia data inside the secure NIH Integrated Data Analysis Platform (NIDAP).

## Author Contributions

GZ, JC, and MJ conceived and designed the project. FL, RH, and GZ developed the pipeline from package creation to deployment. TS and GZ implemented the Shiny application. FL and DS analyzed the data and interpreted the results. SL, LR, and DW provided the dataset and contributed to the discussion of the results. FL and GZ drafted the manuscript and prepared the figures. All authors reviewed and approved the final manuscript.

## Funding

This work was supported in part by NCI Contract No. 75N91019D00024, and the Purdue Institute for Cancer Research and NIH grant P30 CA023168.

## Code Availability

The SPAC workflow code is available at: https://github.com/FNLCR-DMAP/SCSAWorkflow.

## Declarations

The authors have declared that no conflict of interest exists.

---

**Supplementary Fig. 1.**
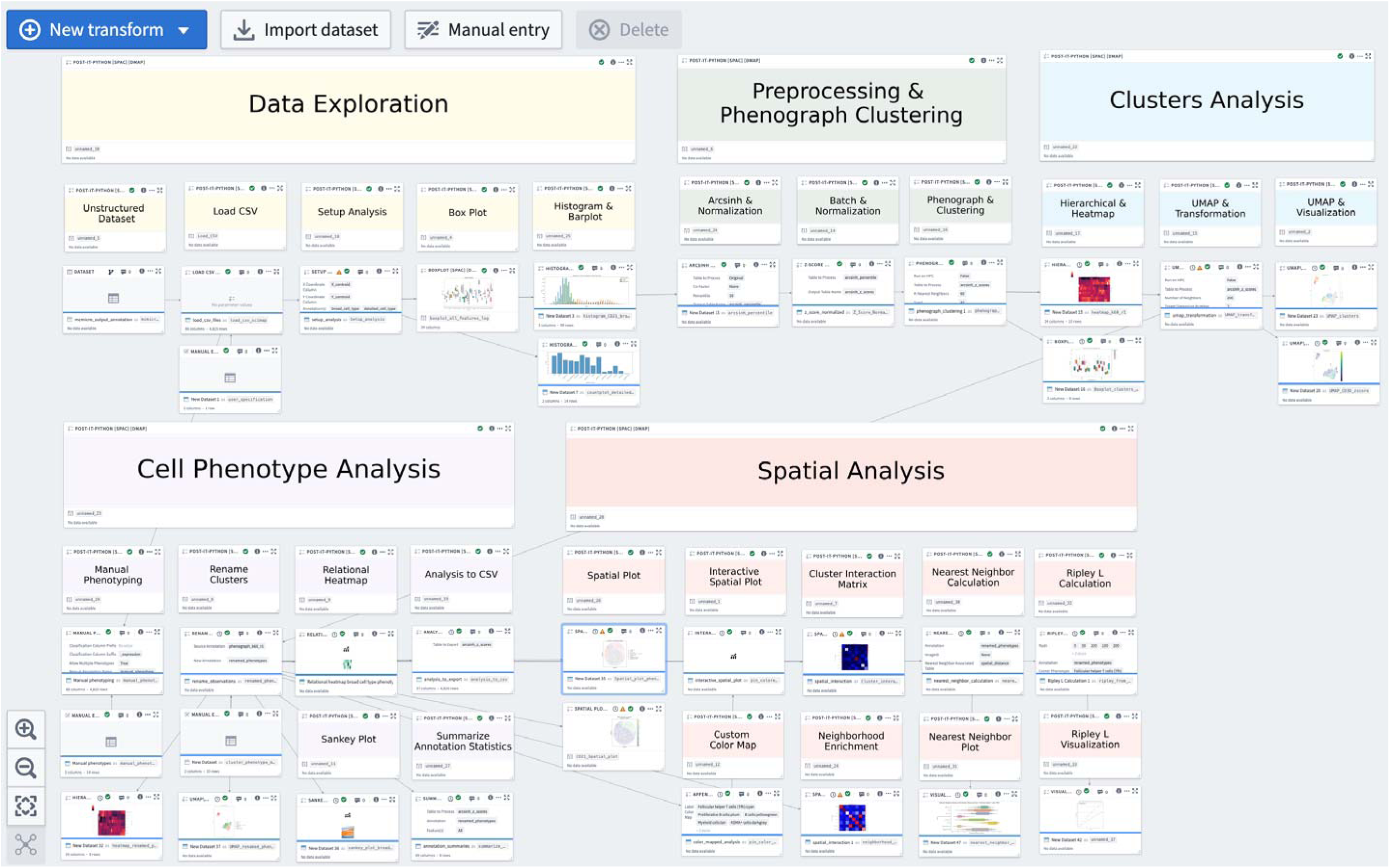
A representative Code Workbook on NIDAP illustrates the SPAC workflow from data aggregation and sampling, exploratory data analysis, feature normalization, clustering, dimensionality reduction, and phenotype annotation to spatial analysis in a unified, modular environment, ensuring consistent data lineage and minimizing format conversion burdens. Each analysis step produces downloadable .csv files and figures ready for scientific presentation. Integrated HPC Connector in PhenoGraph and UMAP modules enable seamless offloading of computations to GPU/CPU resources for efficient processing of large-scale datasets. An example analysis of normal lymph node tissue demonstrates NIDAP’s standardized yet flexible design, enabling bench scientists to configure parameters, launch workflows, and view results in real time without command line expertise, while data scientists can refine robust pipelines, with reproducibility and transparency maintained through version control, parameter tracking, and shared project workspaces.

**Supplementary Fig. 2.**
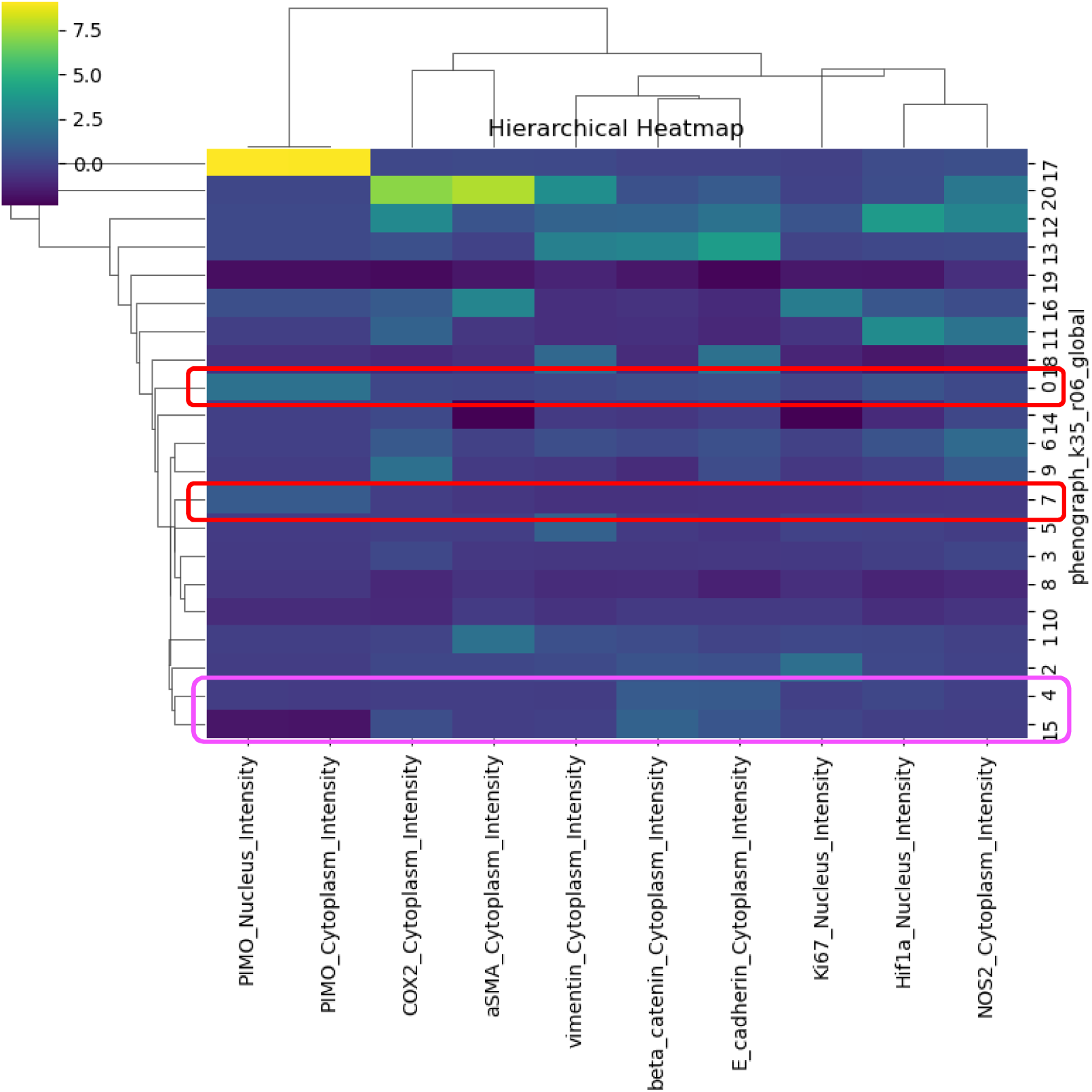
Hierarchical heatmap illustrating the expression profiles of key markers (columns) across PhenoGraph clusters (rows). Each cell represents the z-score of a given marker’s intensity (yellow= higher expression; purple=lower expression). The dendrograms show how clusters and markers group together based on similarity in expression patterns, revealing distinct subpopulations. Clusters 4 and 15, outlined in magenta boxes, exhibit high E-cadherin and β-catenin expression. Clusters 0 and 7, outlined in red boxes, display moderate PIMO expression. These highlighted clusters were subsequently merged and renamed in the final analysis (see Fig. 7).

## Notes

### Competing Interest Statement

The authors have declared no competing interest.

