## Supplemental Figure for "SPAC: a scalable, integrated enterprise platform for end-to-end single cell spatial analysis of multiplexed tissue imaging"

### Slide 1
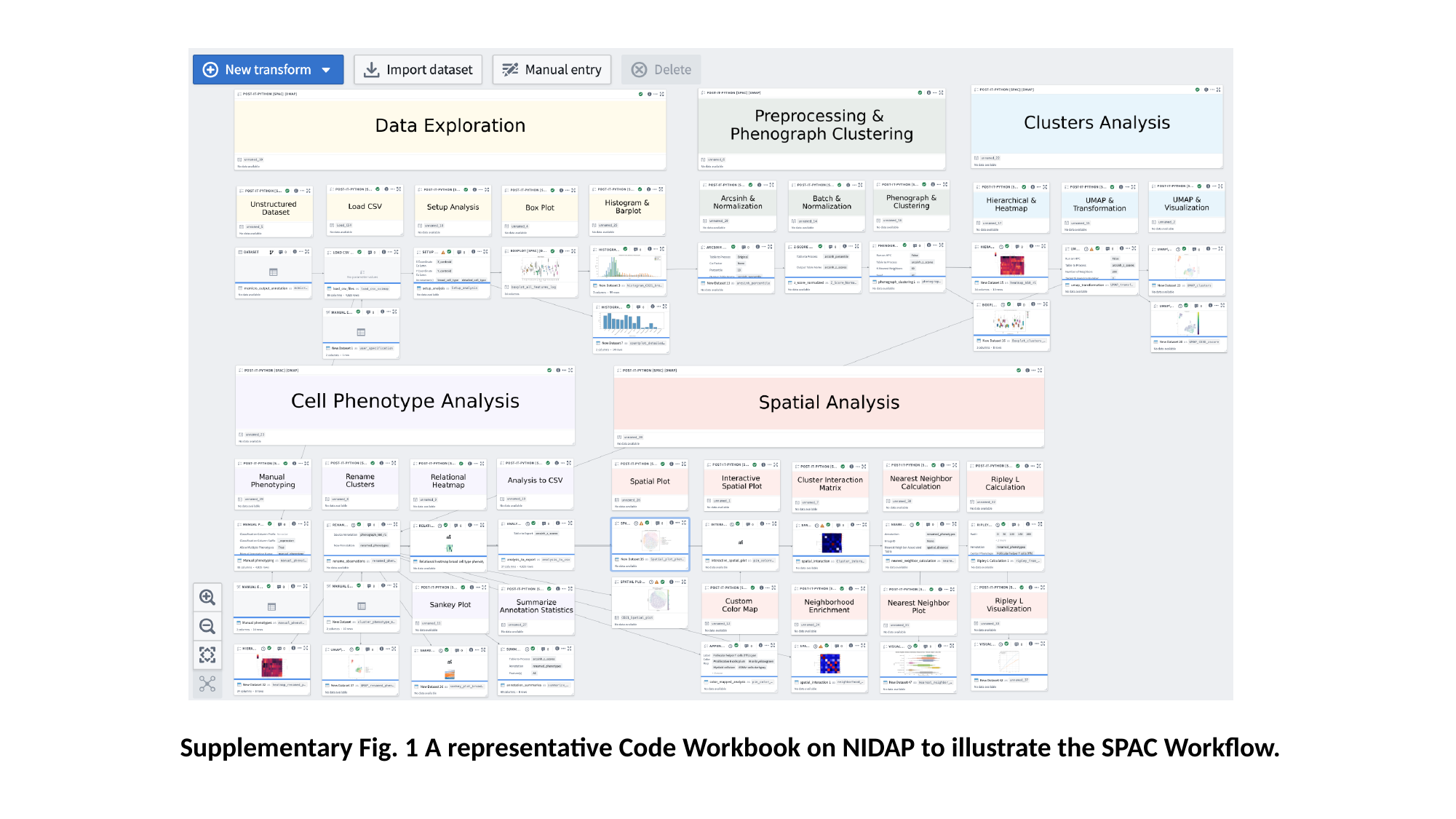

Supplementary Fig. 1 A representative Code Workbook on NIDAP to illustrate the SPAC Workflow.

### Slide 2
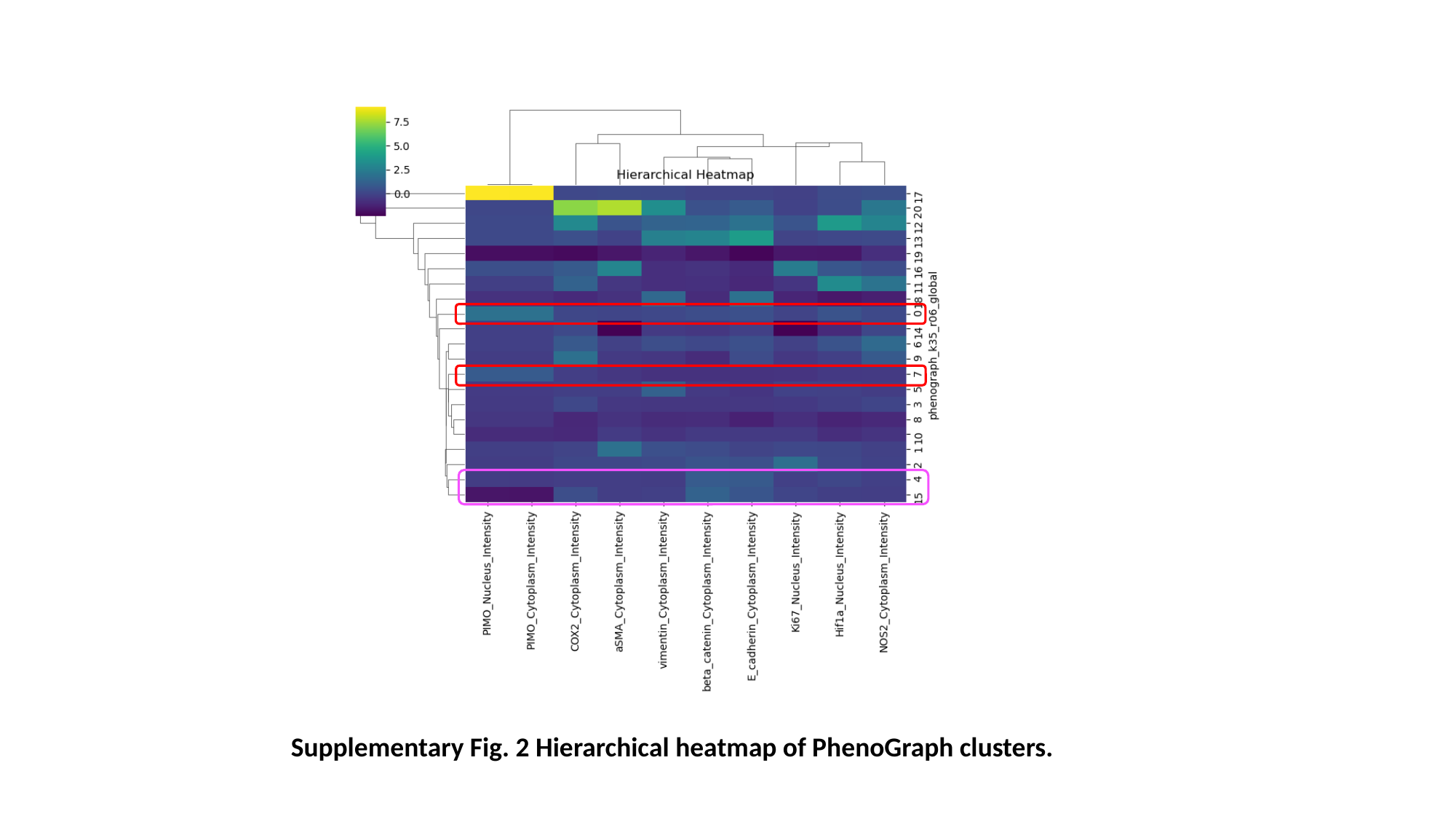

Supplementary Fig. 2 Hierarchical heatmap of PhenoGraph clusters.
